# Divergent demographic strategies drive population dynamics of temperate plants across varying deposition

**DOI:** 10.64898/2026.07.18.739318

**Authors:** Miao Hai-Tao, Ruobing Xue, Yuan Xiaobo, Niu Decao, Shou-Li Li

## Abstract

A central question in biodiversity conservation is whether species can sustain viable population under current and future atmospheric N deposition. Assessing species viability under N deposition requires demographic studies integrating species’ vital rates responses to long-term N deposition across different levels. However, studies of this nature are rare. Our integral projection models (IPMs), parameterized with demographic data, revealed differing responses of two functionally similar coexisting species, *Stipa bungeana* and *Leymus secalinus*, to 12 years of N deposition at low N addition levels (1.15 and 2.30 g N m^−2^ yr^−1^) and high N addition levels (4.60, 9.20, and 13.80 g N m^−2^ yr^−1^) on the Loess Plateau grasslands. We found that the reduced survival across N addition levels was partially compensated by increased contributions from growth, shrinkage, and fecundity, alleviating the population decline of *S. bungeana* (with a longer lifespan and generation time) under different N additions. Contrasting, more positive correlations among vital rate enabled the population of *L. secalinus* (with a shorter lifespan and generation time) to track N additions, with population growth under low N additions and population decline under high N additions. Our results illustrate that the demographic response to N deposition may vary considerably between functionally similar coexisting species, and species with demographic compensation can buffer populations against N deposition while with demographic lability enable populations to track N deposition. Furthermore, our study demonstrates the potential of using life-history traits to predict species’ viability under N deposition, thereby informing biodiversity conservation under global change.

## INTRODUCTION

Global nitrogen (hereafter, N) deposition poses an increasing threat to biodiversity worldwide by altering organisms’ environments (Galloway *et al*. 2008). Thus, predicting local extinction risk and developing proactive conservation strategies for vulnerable species has become increasingly urgency. Whether species can persist under N deposition depends on their ability to maintain viable populations under such novel environmental conditions (Gotelli & Ellison 2002; Issaka *et al*. 2023). However, quantitively assessing the extent to which N deposition can shape population viability remains challenging, because it requires demographic studies that integrate vital rates (e.g., survival, growth, and reproduction) over an entire life cycle in response to N deposition. The impacts of N deposition on individual performance—such as changes in physiology (e.g., leaf N), morphology (e.g., plant height), phenology (e.g., flowering time), and demographic rates—have been extensively studied (Tilman 1987; Sefcik *et al*. 2007; Xia & Wan 2013; Kubert *et al*. 2021). However, we still lack mechanistic insights into the impacts of such changes on long-term population viability. This knowledge gap hinders our capability to strategically manage populations of conservation concern under global N deposition.

Evaluating population viability responses to N deposition requires multi-species demographic studies across a realistic gradient of N addition. Given that N is the most widely limiting nutrient for plant demography (Vitousek & Howarth 1991; Ågren *et al*. 2012), low levels of N deposition may enhance plant performance by alleviating N limitation (Tilman 1987; Phoenix *et al*. 2012), whereas high levels of N deposition can exert detrimental effects through nutrient leaching and mobilization of toxic elements, driven by soil eutrophication and acidification (Spink & Parsons 1995; Högberg *et al*. 2007; Band *et al*. 2022). Currently, most demographic studies have focused on population response to N addition rates (≥10 g N m^−2^ yr^−1^) (Hartnett 1993; Dalgleish *et al*. 2008; Russell & Houseman 2019) that substantially exceed projected future N deposition scenarios (Lamarque *et al*. 2005). Although some demographic studies have examine the demographic response to N addition gradients (Redbo-Torstensson 1994; Smith *et al*. 1999; Gotelli & Ellison 2002; Issaka *et al*. 2023), these treatments did not calibrated against local ambient nitrogen deposition rates (but see Zettlemoyer 2022). Consequently, demographic responses of plant populations to realistic and projected N addition levels remain insufficiently understood. Moreover, such demographic studies have often focused on a single species (Hartnett 1993; Redbo-Torstensson 1994; Gotelli & Ellison 2002) because of the substantial labor and time required for demographic data collection, making it difficult to compare findings and draw robust, generalizable conclusions (Miao *et al*. 2025). Importantly, coexisting species may respond asynchronously to elevated N deposition (Loreau & Hector 2001; Quinn Thomas *et al*. 2010; Hautier *et al*. 2014), with declines in forbs and legumes and increases in grasses due to differences in competitive ability (Issaka *et al*. 2023), highlighting the need to consider multi-species responses for accurate population projections. Therefore, a more comprehensive understanding of how, and to what extent, N deposition regulates population viability is essential for projecting population dynamics under current and future global change scenarios. However, it remains unknown whether functionally similar species inhabiting the same habitats will show similar demographic responses to N deposition.

The direct effects of N deposition on any aspect of individual performance do not directly scale up into impacts on population viability. N deposition may affect vital rates not only in magnitude but also in direction (Gotelli & Ellison 2002; Sefcik *et al*. 2007; Gornish 2014). Furthermore, the extent to which population persistence is affected by N deposition depends on not only the magnitude of N-induced changes in vital rates but also on the sensitivity or elasticity of population growth to these changes (de Kroon *et al*. 1986; Benton & Grant 1999). Because elasticities of vital rates are associated with local selection pressures (Benton & Grant 1996; Caswell 2001), nitrogen deposition–induced changes in elasticity may reshape the strength and direction of selection, thereby influencing optimal life-history strategies. Consequently, assessments of population viability require comprehensive modeling approaches that account for the cumulative effects of N deposition on vital rates and population dynamics.

Here, we examine the demographic response of two dominant plant species, *Stipa bungeana* Trin and *Leymus secalinus* (Georgi) Tzvel, to 12 years of N deposition at different levels in a semi-arid grassland on the Loess Plateau. Atmospheric N deposition in this region has increased substantially in recent decades, driven by socio-economic development in western China (Wei *et al*. 2011; Liu *et al*. 2017). Our study sites have received a N addition starting in 2009 (Niu *et al*. 2018). To quantifying the response of both species to N deposition, we parameterized integral projection models (IPMs) with demographic data collected in 2021 and 2022 (Easterling *et al*. 2000). We used these models to test the following hypotheses: (1) low N deposition positively affects individual performance and subsequently will promote the long-term population growth rate, because of alleviation in N limitation for plant species; (2) high N deposition detrimentally affects individual performance and subsequently will reduce the long-term population growth rate by disrupting nutritional balance; (3) functionally similar coexisting species adopts different life-history strategies in response to N deposition due to differences in competitive ability. If these hypotheses are supported, it can help to inform community-wide management in the face of N deposition.

## MATERIALS AND METHODS

### Study area and experimental design

The study was conducted in a grassland at the Semiarid Climate and Environment Observation of Lanzhou University (SACOL), Gansu Province, China (37°36′ N, 101°19′ E; 1966 m above sea level [asl]). The study area is located on the western Loess Plateau within the arid and semiarid regions (Figure 1). The climate is characterized by a short, dry winter and a long, cool summer. The mean annual temperature is 6.7□, with highest monthly temperature occurring in July (18.9□) and the lowest monthly temperature in January (−7.7□). The mean annual precipitation is 381.8 mm, with >90% occurring as rain during the growing season (≥5□), from April to October (Huang *et al*. 2008). The site is a typical temperate semi-arid steppe, and the vegetation is dominated by perennial herbaceous plants such as *S. bungeana, L. secalinus*, and *Cleistogenes squarrosa* (Niu *et al*. 2018). The study site was fenced to avoid animal disturbance.

**Figure 1.**
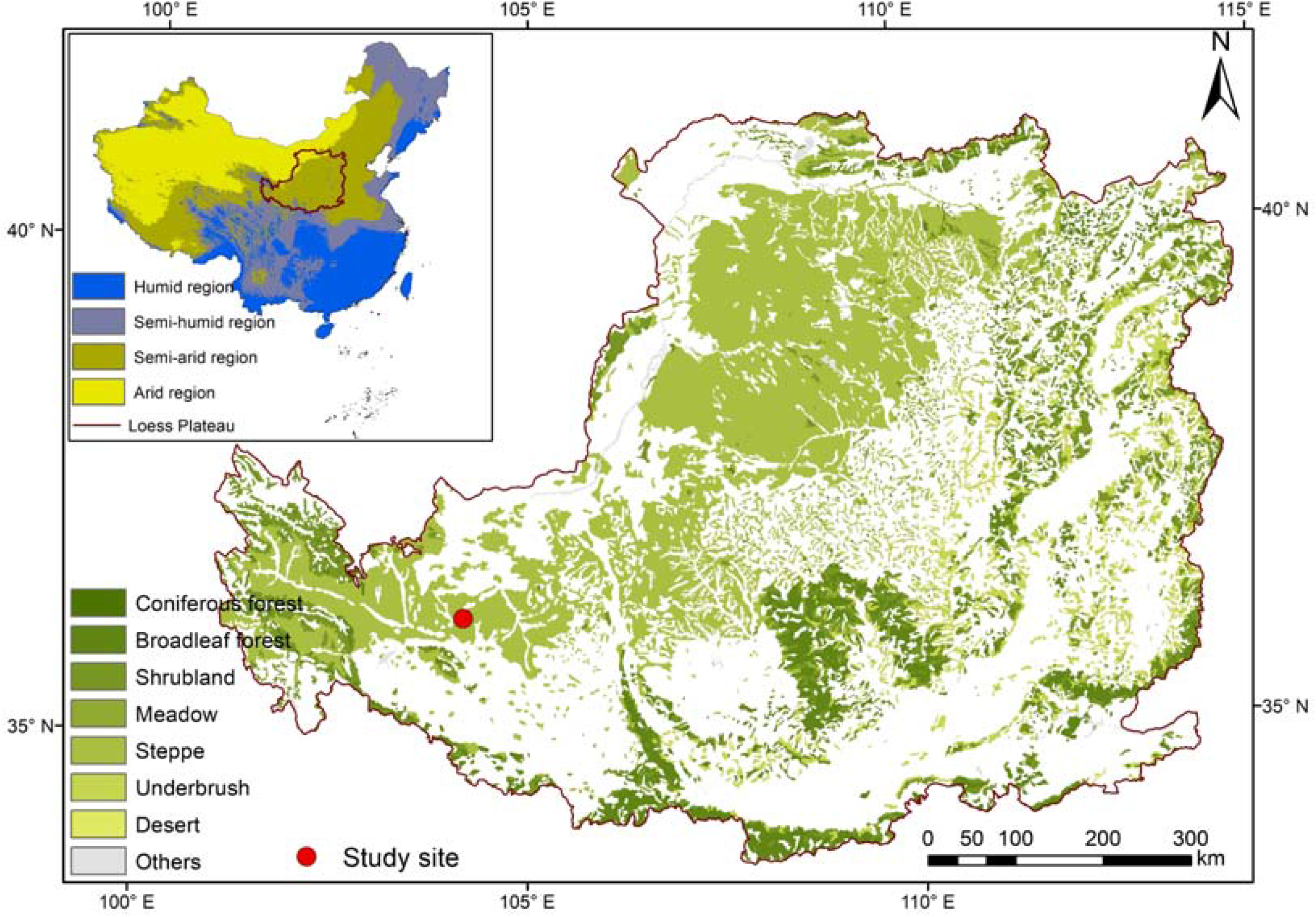
Geographic location of the study site on the Loess Plateau, located in the arid and semi-arid regions of northern China.

A N addition manipulation experiment was established in April 2009 using a completely randomized design. The experiment consisted of six treatments: an ambient condition and five N addition levels (1.15, 2.30, 4.60, 9.20, and 13.80 g N m^−2^ yr^−1^), each with five plots. Each plot measured 1 × 1 m, and was randomly distributed on the landscape with a spacing of approximately 10 m between plots. Hereafter, these treatments are referred to in abbreviated form based on the amount of N added (N0, N1.15, N2.30, N4.60, N9.20, and N13.80). Urea [CO(NH_2_)_2_] was applied starting in May 2009 and repeated twice annually, with 50% of the annual N addition in late May and 50% in late June. The fertilizer was dissolved in 10 L of purified water and evenly sprayed over each plot on a rainy day using a portable sprayer. Loess Plateau is currently experiencing exacerbated atmospheric N deposition (Lü & Tian 2007; Lu *et al*. 2011). To simulate N enrichment at levels comparable to current atmospheric N deposition, the low N addition levels (1.15 and 2.30 g N m^−2^ yr^−1^) were chosen, consistent with the actual deposition rates of 0.2–2.20 g N m^−2^ yr^−1^ at our study site (Lü & Tian 2007). The high N addition levels (4.60, 9.20, and 13.80 g N m^−2^ yr^−1^) were set to simulate N enrichment at levels expected for future atmospheric N deposition. In addition, 10 L of purified water without N was applied to mimic the water effect in the ambient plots.

### Study species and demographic data

To examine the effects of N deposition on the population performance of functionally similar grasses inhabiting the same habitats, we selected two co-occurring temperate species, *S. bungeana* and *L. secalinus*. Both species are perennial graminoids, but differ in life form and reproductive strategy. *S. bungeana* is a bunchgrass with clumped basal shoots that reproduces mainly by seed, whereas *L. secalinus* is a rhizomatous grass with long belowground rhizomes, reproducing primarily through vegetative propagation (Li *et al*. 2022). The two species share similar seasonal growth cycles: both produce new shoots annually, regrow in March–May, flowering occurs in June–July, and set fruit in August–September. The two species are widely distributed on the grasslands of the Loess Plateau, and serve as important forage species (Hu *et al*. 2013; Ye *et al*. 2015).

To examine the effect of N deposition on demographic processes, annual censuses were conducted in August 2021 and 2022 for both species. In the first census, we measured individual plant height, maximum crown diameter, perpendicular crown diameter for *S. bungeana*, and individual plant height and number of leaves for *L secalinus*. We also recorded the reproductive status of each individual and counted the number of spikes produced by each reproductive individual for *S. bungeana*, while no reproductive individuals were found for *L secalinus*. Upon the first measurement, we tagged each individual with a stainless-steel label, and recorded its Cartesian coordinates with the plots to allow tracking through time. In 2022, we checked the survival of all tagged individuals and remeasured plant size and the aforementioned reproductive characteristics on surviving ones. At that time, we also located and measured new recruits within our plots, including new seedlings for *S. bungeana* and new ramets for *L secalinus*. Because adult individuals of *L secalinus* usually produce numerous ramets and tracing the ‘parent’ of new ramets is not feasible without destructive sampling, we assumed that vegetative reproduction in this species is size-dependent. Specifically, the vegetative offspring produced by an individual was assumed to be linearly related to its size (Verburg *et al*. 1996). Accordingly, the number of recruits produced per individual was estimated by multiplying its size (*x*) by the ratio *n*⁄∑*x*, where *n* is the total number of new ramets in the current year and ∑*x* is the sum of the sizes of all individuals in the previous year within each plot (Li *et al*. 2013; Li *et al*. 2015). In total, we tracked 1,061 individuals of *S. bungeana* and 1,213 individuals of *L secalinus* across all plots.

### Demographic rate estimates

To examine the effects of N deposition on the two species’ demography, we used regression models to relate each vital rate to treatment and/or individual size. Demographic data were successively pooled across plots and treatments and then used to fit regression models for the vital rates that collectively determine population dynamics. For *S. bungeana*, six vital rates were modeled: probability of survival, changes in size (i.e., growth and shrinkage), probability of reproduction, number of spikes, probability of recruitment, and the distribution of recruit sizes. In contrast, for *L secalinus*, four vital rates were modeled: probability of survival, changes in size (i.e., growth and shrinkage), number of recruits produced per size-specific plant, and the distribution of recruit sizes. Probability of survival and reproduction were modeled from the Binomial family with the logit link function. Changes in size and distribution of recruit sizes were modeled from the Gaussian family. Number of spikes were modeled from the negative binomial family with the log link function. Probability of recruitment and number of recruits produced per size-specific plant were modeled from the beta family with the logit link function (Cribari-Neto & Zeileis 2010).

To determine the best proxy for plant size in each study species, we used several size metrics to characterize plant survival, size changes, and reproduction. We estimated plant size using four candidate size metrics for *S. bungeana* and three for *L secalinus* (Table S1). These size metrics were derived from the basic morphological measurements described above, either used directly or in arithmetic combinations. All size metrics were also log-transformed to normalize model residuals and reduce skewness where applicable (Baudraz *et al*. 2025). We used the pooled demographic data pooled to construct mixed-effects models for the size-dependent vital rates, with plot included as a random effect, and assessed the overall metric of model performance using Nakagawa’s *R*^*2*^ (Nakagawa & Schielzeth 2013; Johnson 2014; Nakagawa *et al*. 2017). Nakagawa’s *R*^*2*^ includes marginal 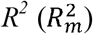 and conditional 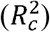 component, where 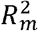 represents the variance explained by the fixed effect and 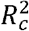 represents the variance explained by the entire model, including the random structure. Because the equations for derive *R*^*2*^ differ depending on the model’s error distribution and the link function (Nakagawa & Schielzeth 2013), we use 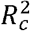to compare metrics in their ability to explain each vital rate. The 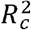 values were normalized prior to averaging them across the different vital rates (Baudraz *et al*. 2025):

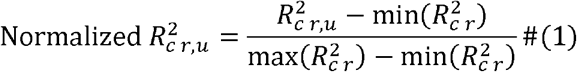

where *r* denotes the vital rate and *u* is the size metric. The variable with the highest mean normalized 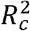 across vital rate models was selected as the proxy for plant size in each study species: log-transformed crone volume for *S. bungeana* and number of leaves for *L secalinus* (Table S2).

To examine how N addition and plant size jointly affect plant vital rates, we developed a set of candidate regression models, including both fixed-effects models and mixed-effects models (Table S3). Fixed-effects models included treatment, plant size, and their interaction as predictors. Mixed-effects models included the same fixed effects, but also incorporated plot as a random effect to account for spatial variation among plots. For the mixed-effects models, we considered different random structures, including random intercepts only, random slopes of plant size, or both random intercepts and slopes, with or without correlations between intercept and slope (Ellner *et al*. 2016; Tredennick *et al*. 2018). Models were fitted using the R packages *nlme* or *glmmTMB* (Bates *et al*. 2015; Brooks *et al*. 2017). We compared candidate models using Akaike’s Information Criterion corrected for small sample size (AICc) to select the best-supported model for each vital rate (Table S4; Burnham & Anderson 2004).

### Population dynamics model

To quantify the effects of N deposition on population dynamics of the two study species, we constructed Integral Population Models (IPMs) for each species based on their demographic processes (Rees & Ellner 2009). IPMs use information on how an individual’s size influences vital rates to project population changes in discrete time (Easterling *et al*. 2000). Each IPM was parameterized with estimated vital rate parameters derived from the best-supported models for each vital rate (Table 1). In our IPMs, the continuous plant size was log-transformed crone volume for *S. bungeana* and number of leaves for *L secalinus*, and the discrete time step (from *t* to *t* + 1) corresponded to one year. The size-structed IPM is

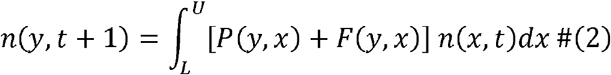

where *L* and *U* are the lower and upper bounds on the range of possible plant size for all treatments. The variable *n*(*x,t*) is the distribution of plant size *x* at time *t*, and n(*y,t* + 1) is the distribution of plant size *y* at time *t* + 1. The expression *P*(*y,x*)+*F*(*y,x*) is called the kernel, *K*(*y,x*), which is a non-negative surface describing all possible demographic transitions from plant size *x* at time *t* to plant size *y* at time *t* + 1. The function *P*(*y,x*) comprises the survival-growth probability of survival of an individual at plant size *x, p*_*s*_(*x*), and the likelihood that component of a IPM and can be decomposed into two functions that determine the the individual will grow from plant size *x* to plant size *y* over a year, *g*(*y,x*), such that P(*y,x*) = *p*_*s*_ (*x*) *g*(*y,x*). The function *F*(*y,x*) differ between the two study species. For *S. bungeana*, the function *F*(*y,x*) comprises sexual reproductive component of a IPM and can be decomposed into four functions that determine the probability of reproduction of an individual at plant size *x, p*_*f*_(*x*), the number of spikes of an individual at plant size *x, f*_ns_(*x*), the probability a recruit establishes *p*_*r*_, and the probability distribution of recruit sizes *f*_*d*_ (*y*), such that *F*(*y,x*) *p*_*f*_(*x*) *f*_*ns*_(*x*) *p*_*r*_ *f*_*d*_(*y*). For *L secalinus*, the function Fcy,xi comprises asexual determine the number of new recruits produced per individual at plant size *x, f*_*nr*_(*x*) reproductive component of a IPM and can be decomposed into two functions that and the probability distribution of recruit sizes *f*_*d*_ (*y*), such that *F*(*y,x*) = *f*_*nr*_(*x*) *f*_*d*_(*y*).

**Table 1.**
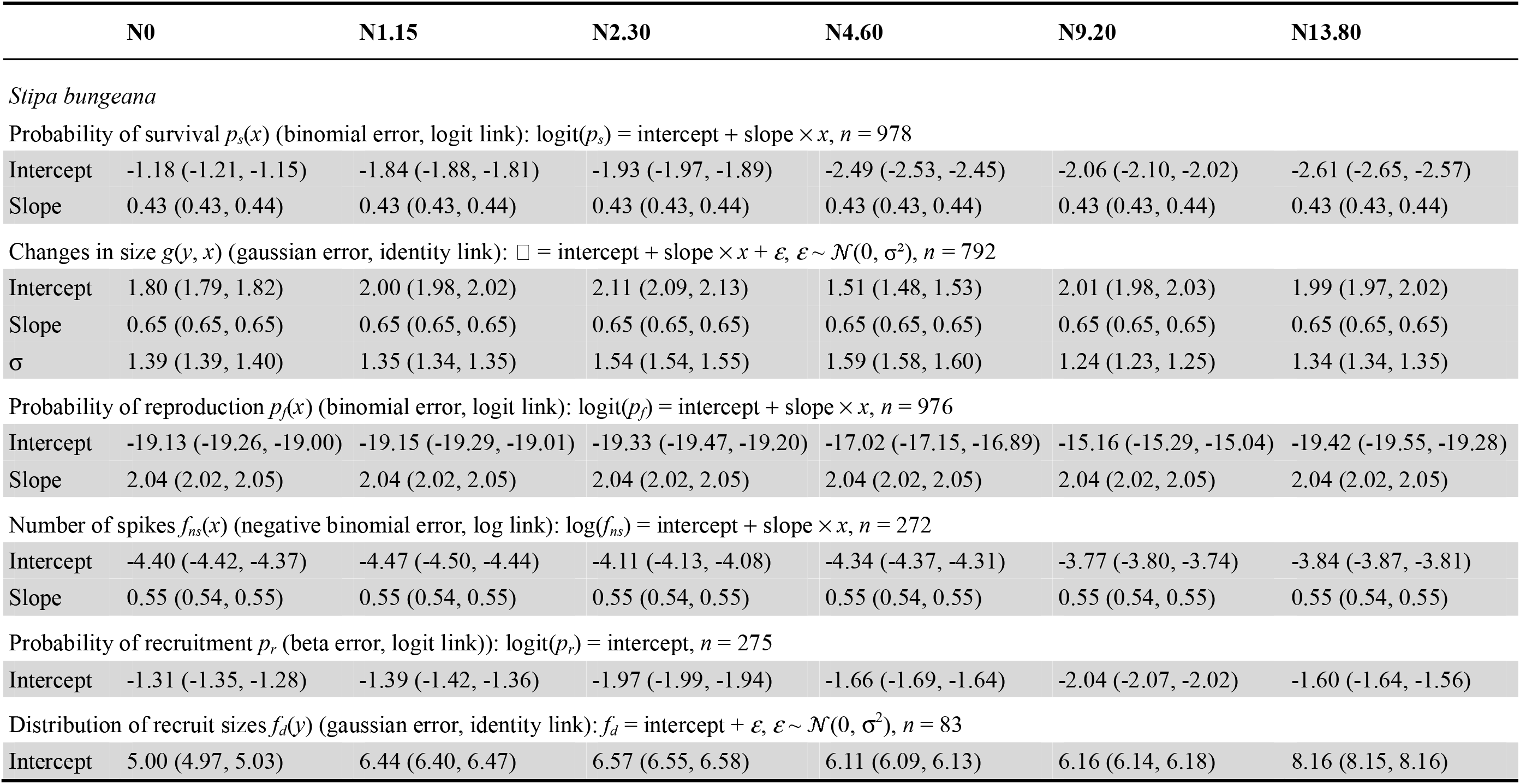
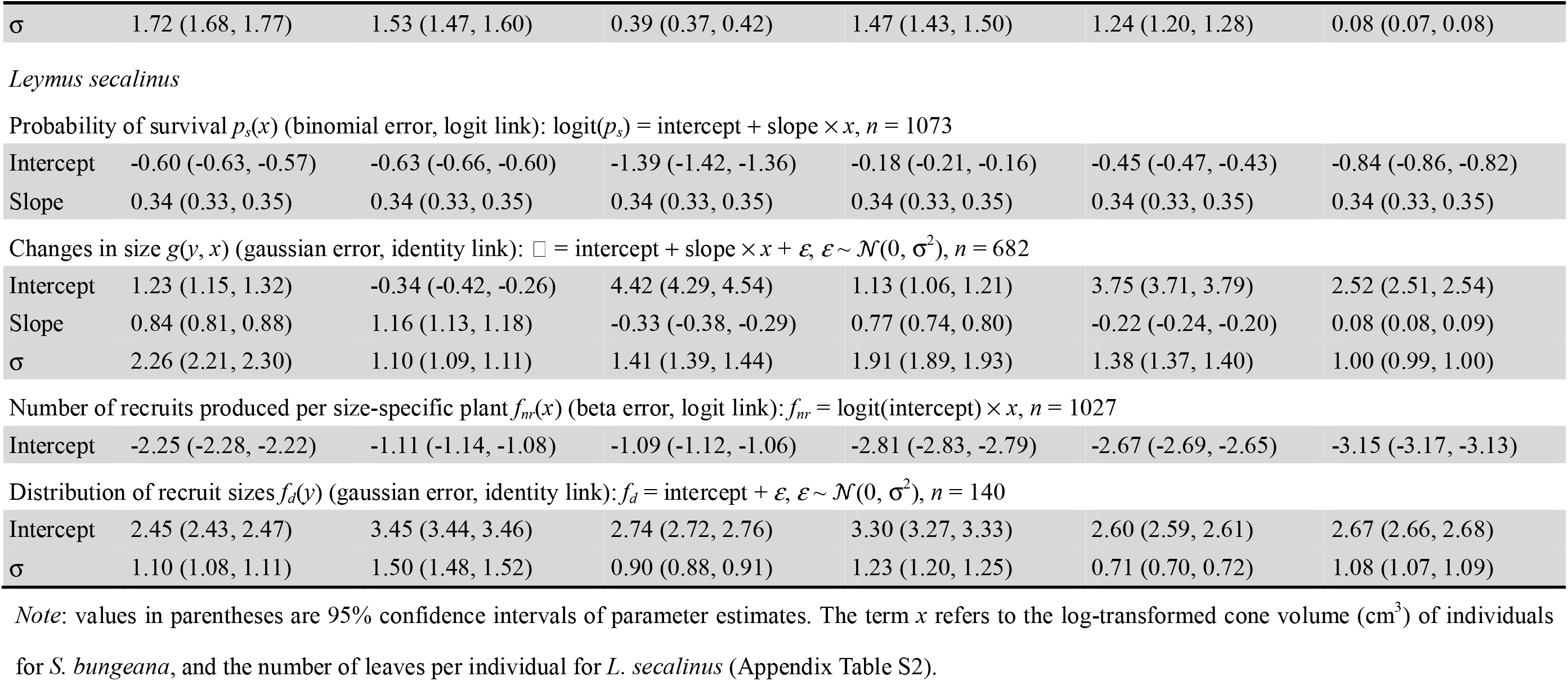
Statistical models and parameter estimates used to construct the kernel for the integral projection model of two co-occurring dominant herbaceous species, *Stipa bungeana* and *Leymus secalinus*, on the Loess Plateau. Best-supported models were chosen with AICc (Appendix Table S4).

To examine the effects of N deposition on long-term population viability, we calculated the population growth rate (λ) of both species under ambient condition and five N addition levels. We discretized the IPM kernel into 200 × 200 matrices using the midpoint rule (Ellner & Rees 2006). The population growth rates, calculated from these matrices as their dominant eigenvalues, indicate whether a population will increase (λ *>* 1) or decline to extinction (λ *<* 1) under conditions of stationary equilibrium (Caswell 1982). To estimate the uncertainty around λ, we bootstrapped the demographic data 1,000 times to obtain 95% confidence intervals (CIs) under ambient condition and five N addition levels for the two study species (Puth *et al*. 2015). Furthermore, we calculated the differences between λ values (Δλ) estimated in each bootstrap to obtain 95% CIs of Δλ between ambient and N additions for both species. In addition, the aforementioned bootstrapped samples were employed in the subsequent population dynamics analysis.

To examine the relative importance of each vital rate on the population growth rate, we conducted prospective perturbation analyses, the elasticity analyses. Elasticity analysis of vital rates quantifies a proportional change in λ in response to a proportional change in vital rate (Zuidema & Franco 2001), which makes comparisons among vital rates (Griffith 2017). By conducting vital rate elasticity analyses, we were able to determine the elasticity values for survival, growth and reproduction separately. Positive elasticity values imply that an increase in a vital rate will increase λ, which is particularly useful for distinguishing the relative importance of positive growth versus shrinkage (Zuidema & Franco 2001). To estimate the uncertainty around vital rate elasticity values, we used the aforementioned bootstrapped samples to obtain their 95% CIs. Additionally, to examine whether the demographic response to N addition may be affected by a plant’s life-history traits, we calculated the generation time and mean lifespan under each N addition level for both species (Ellner *et al*. 2016; Jones *et al*. 2021).

### Life table response experiment

To test how N addition-induced changes in vital rates might have led to differences in λ between ambient and N additions, we conducted retrospective demographic analyses, the life table response experiment (LTRE). An LTRE analysis decomposes the observed difference in population growth rate into the contributions from different vital rates (Caswell 1989). We conducted a fixed-design LTRE on vital rate over a life cycle. The one-factor LTRE analysis follows the formulation

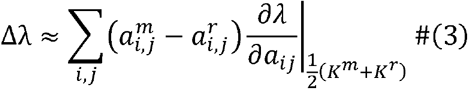

where Δλ is the difference in population growth rate between N addition level *m* and ambient condition *r*. The contribution of each vital rate is calculated by the differences between the vital rate 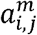 of the large transition matrix *K*^*m*^ under N addition level and the vital rate 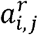 of the large transition matrix *K*^*r*^ under ambient condition multiplied by the sensitivity values of the mid matrix, which is the average of the matrices under N addition level and ambient condition (Caswell 1989). Using the mid matrix reduces deviations arising from the nonlinear nature of vital rate sensitivities (Caswell 1982).

### Demographic strategies

To identify demographic patterns in species’ response to N deposition, we define five distinct demographic strategies regarding the sign and value of vital rates (Figure 2). If both vital rates (e.g., survival and growth) ether increase or decrease with N deposition levels, the population will track either growth or decline according to such demographic additivity (Figure 2a, e, f, j). If vital rates change in the positive directions, the population decline would be buffered by such demographic compensation (Figure 2b-d, g-i). To further quantify the demographic strategies in species’ response to N deposition, we used a randomization approach to evaluate whether the observed data harbored more negative correlation among vital rate contributions than expected by chance (i.e., demographic buffering), or more positive correlation among vital rate contributions than expected by chance (i.e., demographic lability) (Villellas *et al*. 2015). We first calculated the spearman’s correlation coefficient between pairs of vital rate contributions. Second, we estimated the relationship after randomly reassigning all vital rate contributions across N addition levels (repeated 1000 times). Third, we compared the observed numbers of negative correlations (or positive correlations) to the percentiles of the respective null distributions obtained via randomization, and calculated the significance level based on the proportion of values in the respective null distribution.

**Figure 2.**
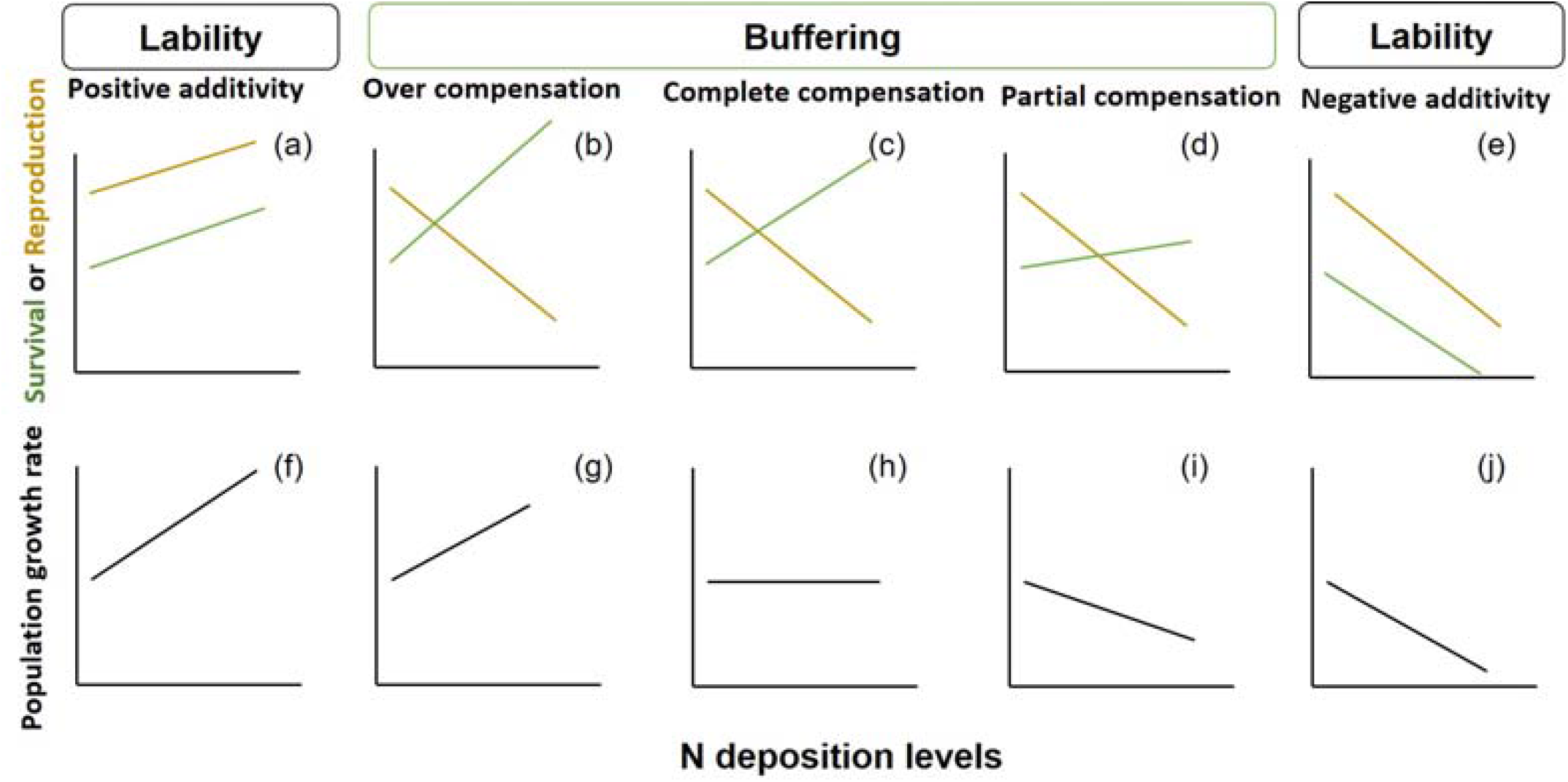
Predictions of how population growth rate (λ) and vital rates (survival probability and reproduction) should vary under N deposition with demographic additivity (a, e, f, j) and demographic compensation (b, c, d, g, h, i). (a, e) Under demographic additivity, both vital rates (lines of different colors) either increase or decrease with N deposition levels, leading to either population growth (f) or decline (j). (b, c, d) Under demographic compensation, vital rates changed in opposite directions, buffering the population decline (g, h, i).

**Figure 2.**
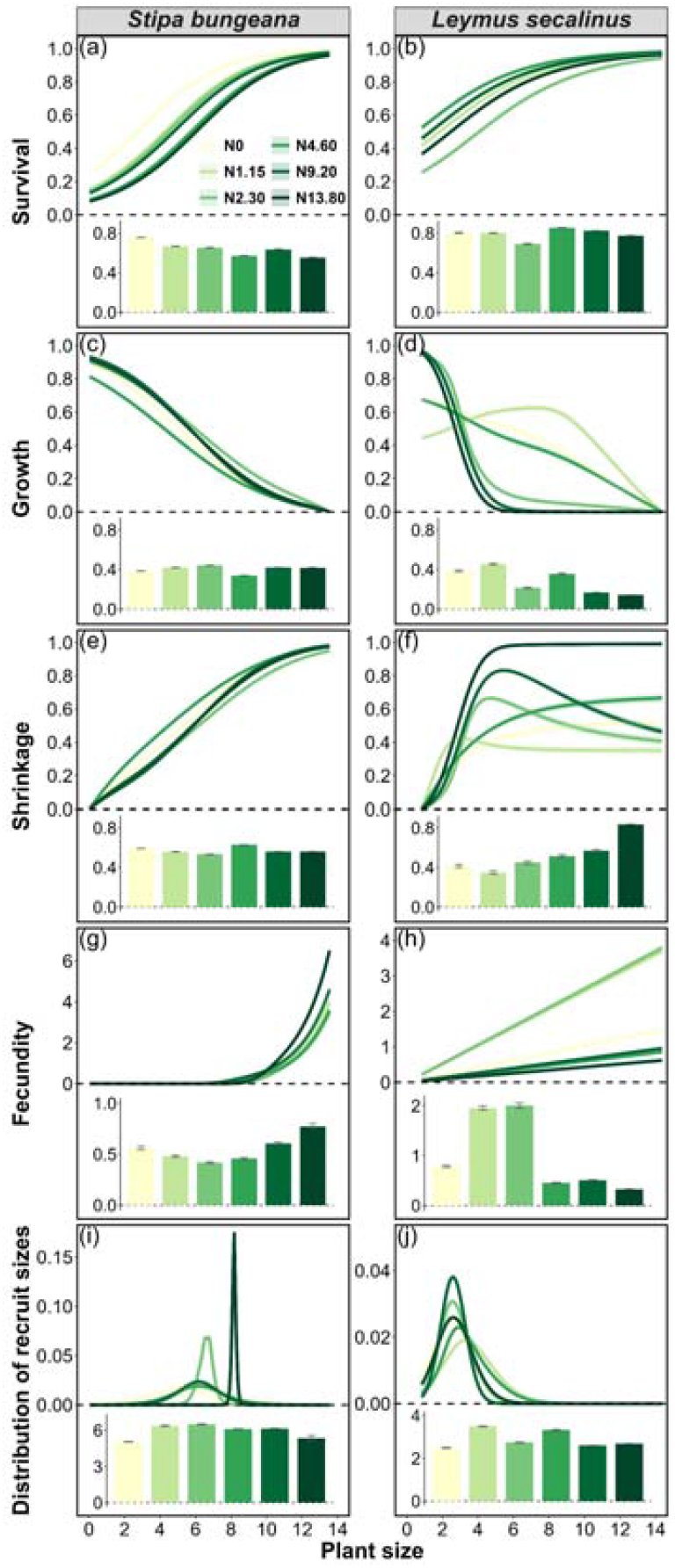
Effects of N addition on the survival (a, b), growth (c, d), shrinkage (e, f), fecundity (g, h), and distribution of recruit sizes (i, j) for two-occurring dominant herbaceous species, *Stipa bungeana* (a, c, e, f, g, i) and *Leymus secalinus*, on the Loess Plateau. Solid lines represent mean values of each vital rate at each size, with 95% confidence intervals in the shaded areas. Mean values of each vital rate across all sizes are given in the bar chart at the bottom of each panel, with error bars representing 95% confidence intervals. Low N addition levels (1.15 and 2.30 g N m^−2^ yr^−1^) were consistent with the actual deposition rates of 0.2–2.20 g N m^−2^ yr^−1^ at our study site, while the high N addition levels (4.60, 9.20, and 13.80 g N m^−2^ yr^−1^) were set to simulate N enrichment at levels expected for future atmospheric N deposition.

All analysis was conducted suing R statistical software v.4.0.5 (R Core Team 2021).

## RESULTS

### The impacts of N deposition on demographic processes

To examine the effects of N deposition on demographic processes of two co-occurring dominant species, *S. bungeana* and *L. secalinus*, we used regression models to relate vital rates to plant size and N addition treatments. We found that the impacts of N addition differed among vital rates, across plant size, and between species (Table 1). The response of survival to N addition was different between the two species. The addition of N consistently reduced the survival of *S. bungeana* across all plant sizes, with a stronger negative effect under high-N addition than under low-N addition (Figure 3a; Figure S1a). In contrast, N addition has divergent effects on the survival of *L. secalinus*: low-N addition consistently decreased its survival across all plant sizes, whereas higher N addition (e.g., N4.60 and N9.20) generally enhanced its survival across all plant sizes (Figure 3b; Figure S1b).

**Figure 3.**
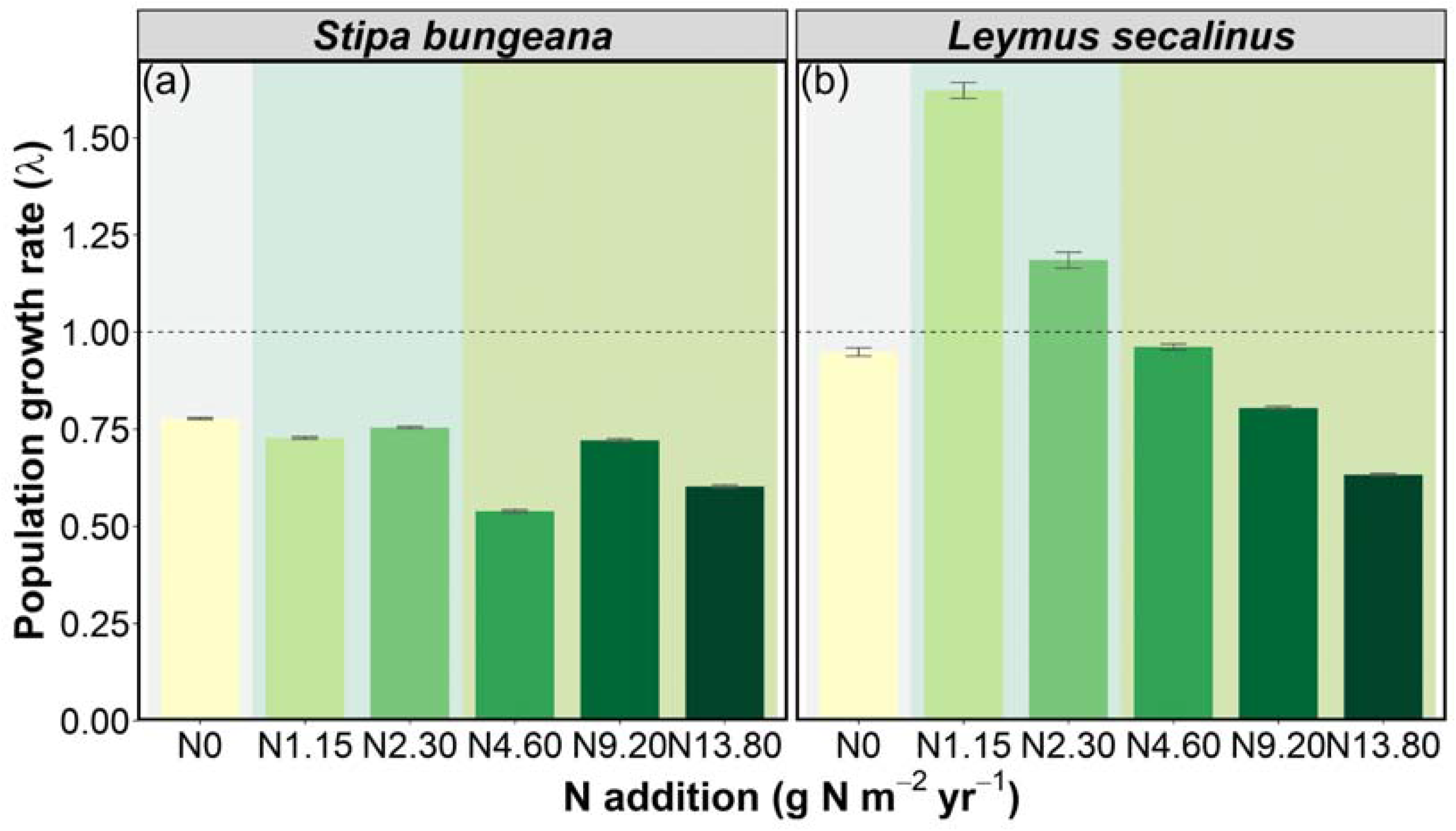
Effects of N addition on the population growth rate of two-occurring dominant herbaceous species, *Stipa bungeana* (a) and *Leymus secalinus* (b), on the Loess Plateau. Error bars indicates 95% confidence intervals. Low N addition levels (1.15 and 2.30 g N m^−2^ yr^−1^; light blue background) were consistent with the actual deposition rates of 0.2–2.20 g N m^−2^ yr^−1^ at our study site, while the high N addition levels (4.60, 9.20, and 13.80 g N m^−2^ yr^−1^; light green background) were set to simulate N enrichment at levels expected for future atmospheric N deposition.

The responses of growth, shrinkage, and fecundity to N addition differed between the two species. Growth generally decreased with plant size in both species, and was generally enhanced by N addition in *S. bungeana* but reduced in *L. secalinus* (Figure 3c, d; Figure S1c, d). In contrast, shrinkage generally increased with plant size in both species, and was generally reduced by N addition in *S. bungeana* but enhanced in *L. secalinus* (Figure 3e, f; Figure S1e, f). Fecundity declined under lower under low N addition but generally increased under high N addition for *S. bungeana*, whereas it increased markedly under low N addition but declined moderately under high N addition for *L. secalinus* (Figure 3g, h; Figure S1g, h). In addition, N addition increased the recruit sizes in both species (Figure 3i, j; Figure S1i, j).

### The impacts of N deposition on population growth rate, generation time, and lifespan

To examine the effects of N deposition on population maintenance, we built IPMs to estimate the population growth rate (λ) for *S. bungeana* and *L. secalinus*. We found that, under ambient condition, λ values were similar for *S. bungeana* (λ = 0.777, 95% CI [0.774, 0.780]; Figure 4a) and *L. secalinus* (λ = 0.949, 95% CI [0.938, 0.959]; Figure 4b). However, the population growth rates were different between the two species under low N addition, with a slight reduction in *S. bungeana* and an increase in *L. secalinus* to above 1 (Figure 4). In contrast, high N addition generally reduced the population growth rates in both species, with the reduction in *S. bungeana* being greater than that under low N addition (Figure 4). The generation time and mean lifespan estimated by the IPMs were longer for *S. bungeana* than for *L. secalinus* under ambient condition: generation time was 6.67□years in *S. bungeana* and 3.32□years in *L. secalinus*; while mean lifespan was 2.88 years in *S. bungeana* and 1.27□years in *L. secalinus* (Appendix S1: Table S5). However, N addition generally shortened the generation time and mean lifespan in both species (Appendix S1: Table S5).

**Figure 4.**
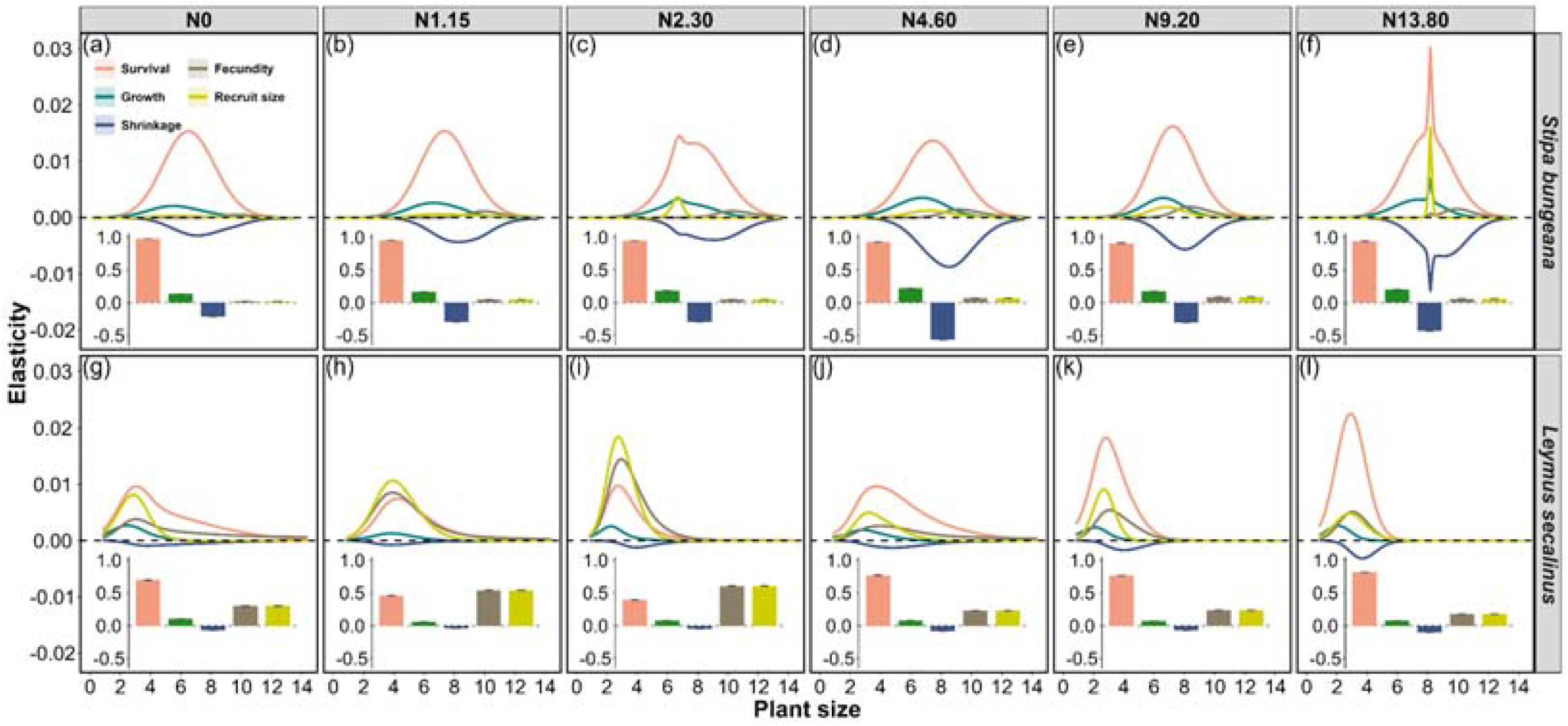
Effects of N addition on the vital rate elasticity of two co-occurring dominant herbaceous species, *Stipa bungeana* (a-f) and *Leymus secalinus* (g-l), on the Loess Plateau. Solid lines represent mean elasticity of each vital rate at each size, with 95% confidence intervals in the shaded areas. Cumulative elasticities of each vital rate across all sizes are given in the bar chart at the bottom of each panel, with error bars representing 95% confidence intervals. Low N addition levels (1.15 and 2.30 g N m^−2^ yr^−1^) were consistent with the actual deposition rates of 0.2–2.20 g N m^−2^ yr^−1^ at our study site, while the high N addition levels (4.60, 9.20, and 13.80 g N m^−2^ yr^−1^) were set to simulate N enrichment at levels expected for future atmospheric N deposition.

### The impacts of N deposition on vital rate elasticity

To identify which vital rates are key to long-term population maintenance under N deposition, we conducted elasticity analyses. We found that the population growth rate was overwhelmingly most elastic to changes in survival for both species under ambient condition (Figure 5a, g). The elasticity to survival in *S. bungeana* declined under N addition, with a stronger reduction under high than low N addition (Figure 5a; Figure S2a). In contrast, the elasticity to survival in *L. secalinus* decreased under low N addition but increased under high N addition (Figure 5g; Figure S2b). In additions, the elasticity of changes in survival increased with smaller plants and decreased with larger individual in both species, peaking at intermediate plant sizes in *S. bungeana* and small plant sizes in *L. secalinus* (Figure 5).

**Figure 5.**
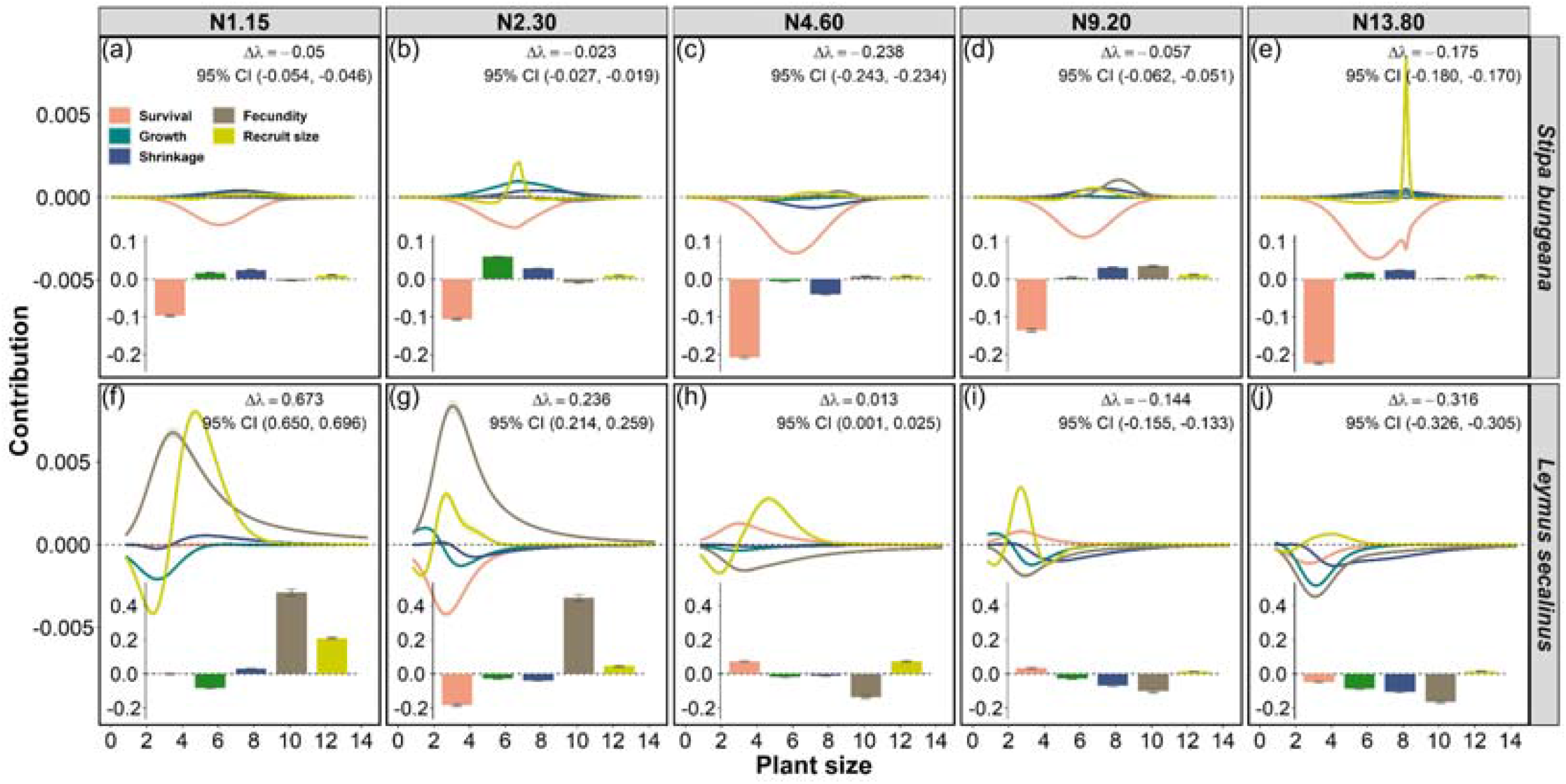
Results of life table response experiment analysis to examine the effects of N addition on the vital rate contribution of two co-occurring dominant herbaceous species, *Stipa bungeana* (a-e) and *Leymus secalinus* (f-j), on the Loess Plateau. Solid lines represent mean contributions of each vital rate at each size, with 95% confidence intervals in the shaded areas. Cumulative contributions of each vital rate across all sizes to the N addition-induced changes in population growth rates (Δλ) are given in the bar chart at the bottom of each panel, with error bars representing 95% confidence intervals. Low N addition levels (1.15 and 2.30 g N m^−2^ yr^−1^) were consistent with the actual deposition rates of 0.2–2.20 g N m^−2^ yr^−1^ at our study site, while the high N addition levels (4.60, 9.20, and 13.80 g N m^−2^ yr^−1^) were set to simulate N enrichment at levels expected for future atmospheric N deposition.

Elasticities to growth, fecundity and recruit size were positive under both ambient and N addition conditions (Figure 5). For *S. bungeana*, these elasticities increased under N addition, with greater increases under high than low N addition (Figure 5b-f; Figure S2c, g, i). In contrast, for *L. secalinus*, the elasticity to growth declined across all N addition levels (Figure 5h-l; Figure S2d), while elasticity to fecundity and recruit size increased under low N addition but declined under high N addition (Figure 5h-l; Figure S2h, j).

Elasticity to shrinkages was negative under ambient and N addition conditions in both species, indicating a negative effect on population maintenance as a result of individuals becoming smaller (Figure 5a-l). This negative effect of shrinkages in *S. bungeana* intensified under N addition, with a stronger increase under high than low N addition (Figure 5b-f; Figure S2e). Conversely, the negative effect of shrinkage in *L. secalinus* weakened under low N addition but strengthened under high N addition (Figure 5h-l; Figure S2f).

### Contributions of vital rates to changes in population growth rates under N deposition

To examine how differences in λ between N addition and ambient conditions were caused by N addition-induced changes in vital rates, we conducted the LTRE analysis. We found that the reduction in λ of *S. bungeana* under N addition was mainly caused by reduced survival of moderate individuals (Figure 6a-e). These negative effects were largely compensated by the positive contributions from growth and shrinkage, resulting in a slight decrease in *S. bungeana* under low N addition (Figure 6a, b). Nevertheless, the additional compensatory increase from fecundity is insufficient to offset the greater decrease in survival under high N addition, leading to a substantial decrease in *S. bungeana* (Figure 6c, d, e). In contrast, the changes in λ of *L. secalinus* under N additions was mainly driven by changes in the fecundity of small plants (Figure 6f-j). Specifically, the increase in λ of *L. secalinus* under low N addition was mainly caused enhanced fecundity, which fully compensated for the reductions in growth or survival (Figure 6f, g). However, the combined negative effects from fecundity, growth, and shrinkage led to the population decline of *L. secalinus* under high N addition (Figure 6h-j).

**Figure 6.**
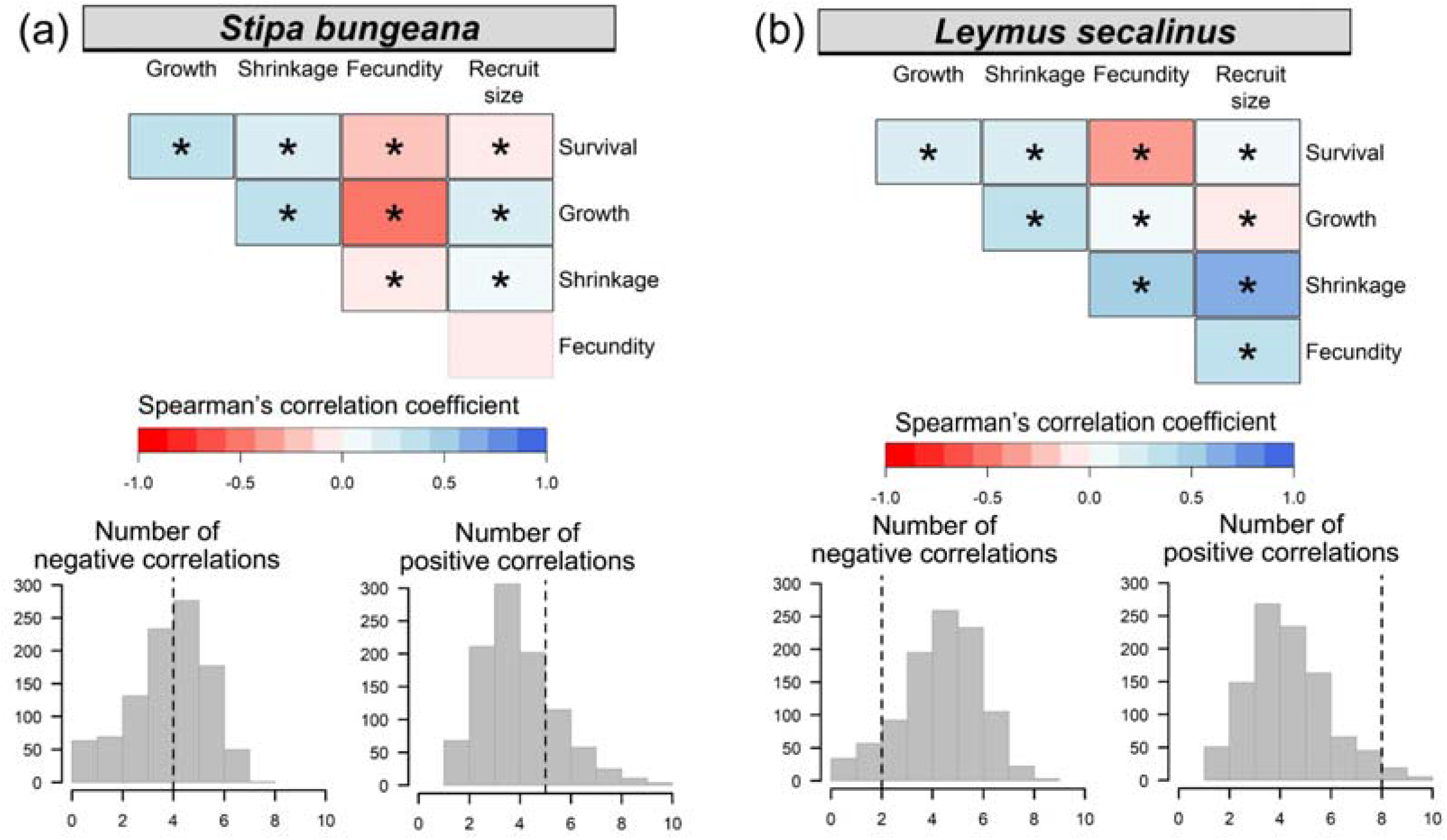
The test of demographic pattern (i.e., demographic compensation or demographic additivity) among vital rates across N addition levels for the two co-occurring dominant herbaceous species, *Stipa bungeana* (a) and *Leymus secalinus* (b), on the Loess Plateau. Correlogram of spearman’s correlation coefficients between the contributions of different vita rates to differences in population growth rates between N addition levels and ambient conditions. Negative correlations are in red, positive correlations are in blue. Boxes with thicker borders and an asterisk indicate significant correlations at *P* < 0.05. The insets show the number of observed negative and positive correlations (vertical dotted lines) compared to the distributions of the number expected by chance obtained through a permutation test (histograms).

To examine the demographic patterns in species’ response to N addition, we used a randomization approach to assess demographic buffering and demographic lability. In *S. bungeana*, we found four significant negative correlations out of 10 possible pairwise correlations across N addition levels, which was not significantly greater than expected by chance (*P* = 0.74; Figure 7a). In contrast, we found eight significant positive correlations out of 10 possible pairwise correlations across N addition in *L. secalinu*, which was marginally greater than expected by chance (*P* = 0.07; Figure 7b). Overall, the population of *S. bungeana* under N addition tend to be more resilient, while the population of *L. secalinu* under N addition is more labile.

## DISCUSSION

### Population dynamics under N deposition

Quantitatively assessing how N deposition might affect the population persistence of natural plants is critical for informing effective biodiversity conservation (Galloway *et al*. 2008; Hautier *et al*. 2014). Comparing the demographic responses of functionally similar species inhabiting the same habitats under elevated N addition can help scientists to evaluate the generality of the findings about the impacts of N deposition on population viability. However, this approach is rarely carried out (but see Dalgleish *et al*. 2008; Zettlemoyer 2022). Here, we examine the demographic responses of two coexisting herbaceous species, *S. bungeana* and *L. secalinus*, to 12 years of different N addition levels on a semi-arid grassland of the Loess Plateau. Although the two species exhibited similar declining population growth rates under ambient conditions, *S. bungeana* is projected to sustain its population decline and *L. secalinus* to thrive under low N deposition. Different demographic responses of functionally similar species to low N deposition have also been documented in diverse systems, including perennial herbaceous plants at the Cedar Creek Natural Area (Gilliam 2006), the extinct and extant confamilial native plants in Michigan prairies (Zettlemoyer 2022), and two rare and endangered fern species in a subtropical forest (Ji *et al*. 2022). Furthermore, such different demographic responses of functionally similar species to elevated N deposition varied not only in direction, but also in magnitude (Gilliam 2006; Zettlemoyer 2022). We observed that decline in population growth rates induced by high N deposition were stronger in *L secalinus* than in *S. bungeana* in our study. Similarity, Zettlemoyer (2022) found that the magnitude of decline in population growth rates differed between extinct and extant confamilial native plants in Michigan prairies. Together, these findings and our results suggests that coexisting species may face contrasting fates under N deposition, resulting in community reassembly, as found in recent studies (Hao *et al*. 2018; Band *et al*. 2022). Therefore, it may not be straightforward to apply demographic information obtained from one species to another, even if they belong to the same functional group. Furthermore, our study demonstrates that conducting demographic studies on multiple co-occurring species is essential for informing community-wide management in face of N deposition.

### Demographic strategies drive the population dynamics under N deposition

Population dynamics under N deposition are largely depended on their demographic strategies in response to such novel environmental conditions. N deposition may affect population viability by unevenly affecting different demographic processes. Demographic compensation among vital rates may allow species to maintain viable populations under N deposition (Villellas *et al*. 2015). In our study, *S. bungeana* exhibited a substantial decrease in survival under N addition, suggesting that the living conditions became less favorable (Zettlemoyer 2022). However, the population successfully compensated for the negative effects of low N addition on survival by enhancing growth and reducing shrinkage, resulting in only a slight decrease in its overall population growth rate. In contrast, such compensatory effects were too small to offset the larger decrease in survival under high N addition, leading to a huge population decline in *S. bungeana*. Demographic compensation under elevated N deposition has been reported in recent studies (Dalgleish *et al*. 2008; Heinken-Smídová & Münzbergová 2012; Gornish 2014; Zettlemoyer 2022). For example, Zettlemoyer (2022) found that increase of growth and flower production in compensation for the reduction of survival alleviated the population decline of *Pycnanthemum tenuifolium*; Heinken-Smídová & Münzbergová (2012) found that the decrease in generative reproduction were adequately compensated by increase in growth from seeding to vegetative adults enabled *Ligularia sibirica* to maintain viable populations under N-rich habitats; Gornish (2014) found that the reduced fecundity of both flowering first-year plants and flowering adults was partially offset by the enhanced survival of flowering first-year plants, forestalling population decline of *Pityopsis aspera*. Together, these findings and our results suggest demographic compensation may help populations to buffer against the negative effects of N deposition on population maintenance.

N deposition may affect population viability by evenly affecting different demographic processes. Positive demographic additivity in response to N deposition may foster population growth, whereas negative demographic additivity may drive population decline (Figure 2). In our study, we observed more positive correlations among vital rates in response to N addition levels for *L secalinus*, suggesting that the population tends to track N addition deposition. In fact, such demographic lability promoted the population growth of *L secalinus* under low N addition, whereas it led to population decline of *L secalinus* under high N addition. Similarly, Redbo-Torstensson (1994) found that low N-deposition induced increased in survival, fecundity, and seedling recruitment promoted the increased the density of *Drosera rotundifolia*; Zettlemoyer (2022) found that the high N-deposition induced decline in survival and flower production accelerated the population decline of *Penstemon digitalis*; Dalgleish et al. (2008) found that decline in survival and production of vegetative tiller under high N addtion reduced the population growth of *Koeleria macrantha*. These findings, together with our results suggest that demographic additivity may lead to different outcome under varying N depositions levels.

The demographic strategies of plant species in response to N deposition may be regulated by life-history traits. Generation time and lifespan are two key life-history traits that may affect plants’ capacity to cope with environmental variation (Garcia *et al*. 2008; Compagnoni *et al*. 2021). Previous demographic studies have demonstrated that plants with short generation time have stronger responses to environmental variability than those with longer generation time (Garcia *et al*. 2008; Compagnoni *et al*. 2021). This pattern may be driven by short-lived species’ higher vital rate variability (Morris *et al*. 2008), which may led population to track rather than buffer environmental fluctuations (Koons *et al*. 2009; Jongejans *et al*. 2010; Koons *et al*. 2014; Compagnoni *et al*. 2021). In our study, the demographic lability in *L secalinus* (with a shorter generation time and lifespan) led it to track N deposition, while the demographic compensation in *S. bungeana* (with a longer generation time and lifespan) allowed it to buffer N deposition. Such distinct demographic strategies arise because fecundity was more variable than survival for the short-lived species (Garcia *et al*. 2008). Indeed, our LTRE results showed that change in fecundity contributed the most to population growth under N deposition for *L secalinus*, while reduction in survival made the largest contribution to population growth under N deposition for *S. bungeana*. Our results, with these similar findings, suggest the key life-history traits like generation time and lifespan might be key predictors of species’ demographic strategies to cope with N deposition and life-history trait-based approaches in population dynamics may help inform biodiversity conservation under N deposition.

### Management implications in the face of N deposition

Livestock grazing is a common land-use activity on the grasslands of the Loess Plateau (Zhang *et al*. 2016), providing vital resources for cattle and sheep (Zhang *et al*. 2017). We found that survival was key for population maintenance in both species, indicating that grazing activities should be regulated to minimizes mortality. Additionally, shrinkage had a considerable negative elasticity on the population growth rate of *S. bungeana* in the current study, suggesting that a reduction in plant size is detrimental to the population maintenance of such longer-lived species. Reduced plant size under heavy grazing have been widely documented on the Loess Plateau grasslands, including declines in aboveground biomass, plant height, and vegetation cover (Gao & Carmel 2020; Xiang *et al*. 2023; Cao *et al*. 2024). These findings together with our results suggest that grazing activities should also be controlled to avoid large increases in shrinkage. Furthermore, we found that clonal reproduction is more sensitive to population growth rate of *L. secalinus*, and the decrease in clonal reproduction made the largest contribution to changes in population growth rate of this species under N deposition. A meta-analysis based on 114 studies worldwide revealed that moderate livestock grazing can enhance grasses’ asexual reproduction by increase the number of tillers (Wentao *et al*. 2023). Therefore, livestock grazing should also be controlled at a level that can promote asexual reproduction. Given survival, shrinkage, and asexual reproduction are relatively easy to monitor, they could potentially serve as alert signals for managing plant populations of temperate grasslands under N deposition.

Management strategies should be tailored to the species level as much as possible. Our study site has always been fenced to avid animal disturbance. The populations of *S. bungeana* declined under current conditions and different levels of future N deposition, suggesting that the current grazing regime is unsustainable under either present and future N deposition scenarios. Adjusting the grazing regime could help to reverse population decline (Andrello *et al*. 2012; Miao *et al*. 2025). Future studies are needed to determine the optimal grazing regime, in terms of intensity and season, to maintain viable populations of *S. bungeana* in the face of N deposition. In contrast, our findings of *L. secalinus* suggest that the current grazing regime is likely sustainable for this species under both current and low N deposition conditions. However, further high N deposition is expected to lead to population decline of *L. secalinus*, highlighting the need for grazing regimes to be adapted to the specific living conditions in the future.

## Acknowledgements

This work was funded by the National Natural Science Foundation of China (31971423) and the cost-share exchange projects between the National Natural Science Foundation of China and the Royal Society of the United Kingdom (32011530169).

## Author contributions

SLL and DN conceived the study. RX collected the data, HTM constructed models and wrote first draft of the manuscript, and all authors contributed substantially to revisions. All authors approved the final manuscript.

## Data availability

Data and code are archived in Zenodo (xxxx) and are available for peer review via the following private link: xxxx

